# Evolutionary and Functional Lessons from Human-Specific Amino-Acid Substitution Matrices

**DOI:** 10.1101/2020.05.09.086009

**Authors:** Tair Shauli, Nadav Brandes, Michal Linial

## Abstract

The characterization of human genetic variation in coding regions is fundamental to our understanding of protein function, structure, and evolution. Amino-acid (AA) substitution matrices such as BLOSUM (BLOcks SUbstitution Matrix) and PAM (Point Accepted Mutations) encapsulate the stochastic nature of such proteomic variation and are used in studying protein families and evolutionary processes. However, these matrices were constructed from protein sequences spanning long evolutionary distances and are not designed to reflect polymorphism within species. To accurately represent proteomic variation within the human population, we constructed a set of human-centric substitution matrices derived from genetic variations by analyzing the frequencies of >4.8M single nucleotide variants (SNVs). These human-specific matrices expose short-term evolutionary trends at both codon and AA resolution and therefore present an evolutionary perspective that differs from that implicated in the traditional matrices. Specifically, our matrices consider the directionality of variants, and uncover a set of AA pairs that exhibit a strong tendency to substitute in a specific direction. We further demonstrate that the substitution rates of nucleotides only partially determine AA substitution rates. Finally, we investigate AA substitutions in post-translational modification (PTM) and ion-binding sites. We confirm a strong propensity towards conservation of the identity of the AA that participates in such functions. The empirically-derived human-specific substitution matrices expose purifying selection over a range of residue-based protein properties. The new substitution matrices provide a robust baseline for the analysis of protein variations in health and disease. The underlying methodology is available as an open-access to the biomedical community.

## Introduction

The study of population genetic variation is imperative to the understanding of human evolution. Such understanding opens the door to countless scientific and medical applications. Examples include tracing ancient migration patterns, estimating the pathogenicity of genetic variants, identifying functional elements in the genome, and assessing the adaptivity and conservation of genes and other genomic regions. Gaining such insights is predicated on an accurate background model for the dynamics of human genetic variation.

The majority of genetic variants result from replication errors, cytosine de-aminations, and gene conversion (1, 2). Once a new variant is introduced into a population, its spread among individuals is governed by genetic drift and natural selection, which is affected by the evolutionary utility of the variant (3). At any point in time, in a given population, a variant’s abundance is measured by its allele frequency (AF). In the present human population, the vast majority of the observed variants are rare and population-specific (4).

Various approaches model the dynamics of genetic variation from different angles using these principles. One approach uses comparative genomics between human genomes and other related species (5, 6). However, parameters such as demographic history (e.g., migration, admixture, ancestral population-structure, parental age, generation time) differ among mammals, and thus reduce the accuracy of such phylogenetic estimates (6). A different method directly calculates human-specific mutation rates by counting de-novo mutations within parent-child samples. The short timescale of such analyses (up to a few generations) yields a good model for the introduction of new variants, but mostly ignores the effects of natural selection (4).

Genetic-variation models that concern coding regions and proteomic variation are of particular interest, as they aim to reflect the forces of selection that act on protein structure and function. Traditionally, studies of protein evolution make use of amino-acid (AA) substitution matrices such as PAM and BLOSUM (7–9) as models of genetic variation. These matrices score AA substitutions by their likelihood, as derived from empirical observations. Specifically, the construction of these matrices relied on multiple sequence alignments of evolutionarily related protein sequences across species, such as whole protein homologs (10, 11) or highly conserved protein regions (7). BLOSUM and PAM provide sets of related scoring matrices, which aim to capture a range of evolutionary distances. BLOSUM_62_, named for its use of protein blocks with 62% identity or less, is the most prominent AA scoring matrix. It serves as the default choice for BLAST, as well as other commonly used bioinformatics tools (12, 13).

Despite their widespread use, existing AA substitution matrices were not designed to handle proteomic variation within species, and specifically within humans (14, 15). An additional limitation of cross-species AA substitution matrices is the lack of directionality of substitutions (i.e., not distinguishing between substitution of a first AA to a second AA from a substitution of the second to the first). Furthermore, the BLOSUM matrices do not aim to estimate the probabilities of AA substitutions explicitly. Instead, they assign scores. Finally, both BLOSUM and PAM model substitutions at the AA resolution, whereas codon resolution may be more useful for portraying short evolutionary distances such as within the modern human population (16, 17).

In this study, we provide a probabilistic model of codon substitutions and AA substitutions derived from genetic variation observed in the human population. To this end, we exploit a rich collection of ~7M polymorphic sites in the exomes of over 60,000 unrelated, healthy individuals extracted from the Exome Aggregation Consortium (ExAC) (18). From this comprehensive dataset, we constructed a set of human-specific substitution matrices, which offers a baseline for studying genetic and proteomic variation and drawing evolutionary and functional insights. We examine the information contained in our novel matrices and compare them to the currently used matrices. We further used this methodology to expose protein functional constraints, as reflected by post-translational modification (PTM) sites and the residues that coordinate ion-binding in proteins. We provide the community with a generic framework that utilizes aggregated genetic data to produce substitution matrices for a broad range of taxonomic and evolutionary contexts.

## Results

### Constructing human-specific coding substitution matrices

To construct an amino-acid (AA) substitution matrix that is specific to the human population (Fig.1), we merged data from two complementary sources: (i) the ExAC population database, which reports >7M high-quality single nucleotide variations (SNVs) from exome sequences of 60,706 non-related humans (18); (ii) genomic annotation for all human coding genes. By projecting the ~7M SNVs on top of the gene annotations, we inferred 4.8M observed codon substitutions, found in 37% of all codons in the human exome. Overall, 33% of the substitutions are synonymous, 2% are stop-gain, and 65% are missense.

From these codon substitutions and their allele frequencies (AFs), we constructed a 61×61 codon substitution matrix, denoted *HC1* (standing for Human Codon substitutions). Rows and columns in the matrix represent all coding codons (i.e., excluding the three stop codons), with rows representing the source codons and columns representing the target codons. Rows are normalized so that each entry represents the conditional probability of the codon substitution.

To provide substitution probabilities at the AA resolution, we transformed *HC*^*1*^ into a 20×20 amino-acid (AA) substitution matrix, denoted *HA*^*1*^ (i.e., Human Amino-acid substitutions) (19). However, most codon substitutions cannot occur by substituting a single nucleotide. For example, substituting glycine (coded by GGN) to proline (coded by CCN) would require the mutation of at least two nucleotides. Therefore, *HC*^*1*^ is a sparse matrix, in which 84% of all codon pairs have zero probabilities. Likewise, *HA*^*1*^ contains non-zero values in only 42.5% of all AA pairs. To model all substitutions, we considered three consecutive transitions of *HC*^*1*^, thereby deriving a complete codon substitution matrix *HC*^*3*^ (Fig. 1*B*). We also transformed *HC*^*3*^ into an amino-acid substitution matrix, *HA*^*3*^ (Fig. 1*C*). The numeric values of the four matrices, *HC*^*1*^, *HA*^*1*^, *HC*^*3*^, and *HA*^*3*^, are provided in Supplementary Dataset S1.

**Figure 1.**
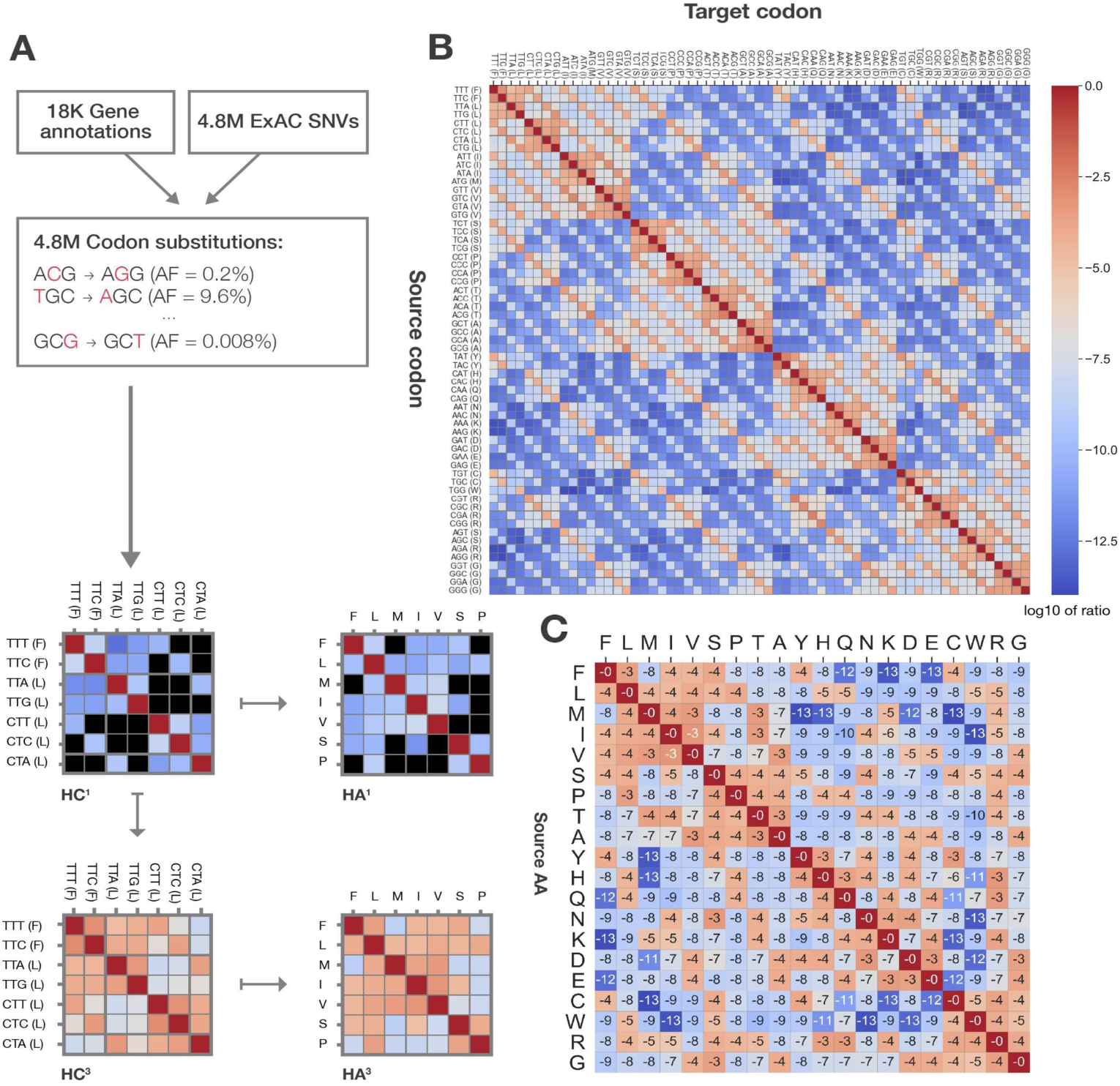
General outline of pipeline and it’s outputs. (A) We combined 4.8M SNVs (from the ExAC database) with 18K gene annotations (from GENCODE) to construct a human-specific codon substitution matrix, denoted *HC*^*1*^, where each entry represents a codon substitution probability. To also represent codon substitutions that differ by more than a single nucleotide, we extended the sparse *HC*^*1*^ into a complete *HC*^*3*^ matrix through a Markovian process. We further consider the two matrices at amino-acid rather than codon resolution, deriving *HA*^*1*^ and *HA*^*3*^, in which each entry represents the probability of an AA substitution. (B) The values of *HC*^*3*^. (C) The values of *HA*^*3*^ (numbers are in log_10_ scale).

### Characteristics of amino-acid substitutions in the human population

We questioned whether the observed 20×20 AA substitution frequencies are exclusively determined by the signal reflected by the 4×4 single-nucleotide substitutions. To this end, we constructed a 4×4 single-nucleotide mutation matrix, denoted *HN*^*1*^, developed from all the reported SNP variants (Fig. 2*A*, Supplementary Dataset S2). This matrix accounts for all 16 single-nucleotide mutation frequencies observed in coding regions across the human population. From *HN*^*1*^, we derived a 20×20 AA substitution matrix, denoted *HAN*^*1*^ (i.e., Human Amino-acid Nucleotide substitutions), which aim to represent the expected AA substitution probabilities under the background model of single-nucleotide substitution propensities (Supplementary Dataset S2). We then compared the expected values of *HAN*^*1*^ to the empirically observed values of *HA*^*1*^ through an element-wise division of the two matrices (Fig. 2*B*). Substitutions of tryptophan (W) to cysteine (C) or serine (S) are roughly 8-fold lower than expected by this naive nucleotide-based background model. Similarly, a substitution from isoleucine (I) to lysine (K) is 16-fold lower than expected. Overall, we reveal that many of the expected-to-observed ratios are far from one (11% are >2 or <0.5, and 20% are >1.5 or <0.66), reflecting the prominence of natural selection even at the resolution of single amino-acids.

**Figure 2.**
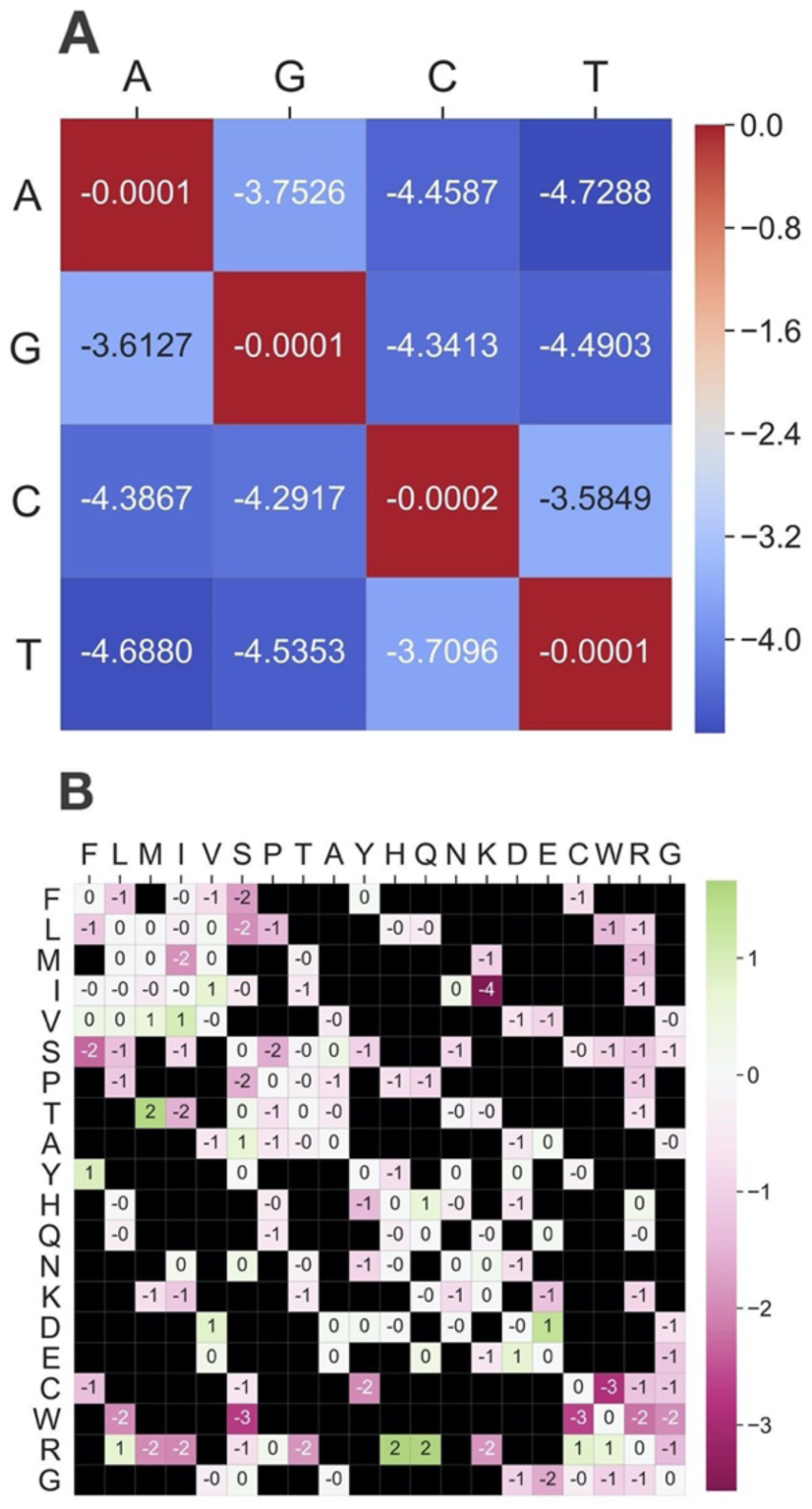
The properties of human-specific amino-acid substitutions (A) The values of *HN*^*1*^, the human-specific nucleotide substitution matrix (B) Expected-to-observed ratios of *HA*^*1*^entries. The 4×4 frequencies of single-nucleotide substitutions determine the expected background model. A ratio close to 1 signifies a substitution whose frequency is mostly determined by the nucleotide content of the codons. Ratios lower than 1 (colored in pink) represent substitutions that are positively selected to be more common than would be expected by their nucleotide compositions. Likewise, ratios higher than 1 (colored green) represent negative selection.

An essential feature of a probabilistic substitution model is the notion of directionality. We consider directionality as the potential asymmetry between the substitution of the first AA to a second AA vs. the opposite substitution of the second to the first. The traditional taxonomy-based matrices (e.g., BLOSUM and PAM) are scoring matrices, which are symmetric by design, and thus lack such directionality. Our substitution models are transition matrices, derived from a single taxon. These properties allow the polarization of the variants using allele frequency information so that the opportunity to explore such asymmetry presents itself. The ratio between each AA substitution to its opposite can be used to measure such asymmetry (Fig. 3*A*). The strongest directionality is associated with tryptophan (W), which shows a much higher tendency to be substituted into than to be substituted by other AAs. Tryptophan (W) is a biochemically distinct amino acid. It is associated with minimal flexibility, maximal hydrophobicity, and bulkiness. It has the most considerable interface propensity and the lowest solvation potential (20). It has also been implicated in the pathogenicity (21). We highlight extreme signals of asymmetry, which we define as substitution ratios higher than 5 (Fig. 3*B*). This representation elucidates patterns of strongly directional substitutions. In addition to tryptophan (W), valine (V) and isoleucine (I) also stand out as hubs of asymmetry, albeit with reversed directionality. That is, whereas tryptophan emerges as a target hub, valine and isoleucine tend to be the source of substitutions. Several directional signals are driven by the substitution preferences of specific codons of the substituted AA. For instance, we observe that tyrosine (Y) is 6 times more likely to substitute valine (V), which is determined by the codon substitution frequency of GTT (V) and GTA (V) to TAT (Y). We observe a weaker opposite tendency for GTA (V), GTG (V), and GTC (V) to TAC (Y) (Supplementary Fig. S2).

**Figure 3.**
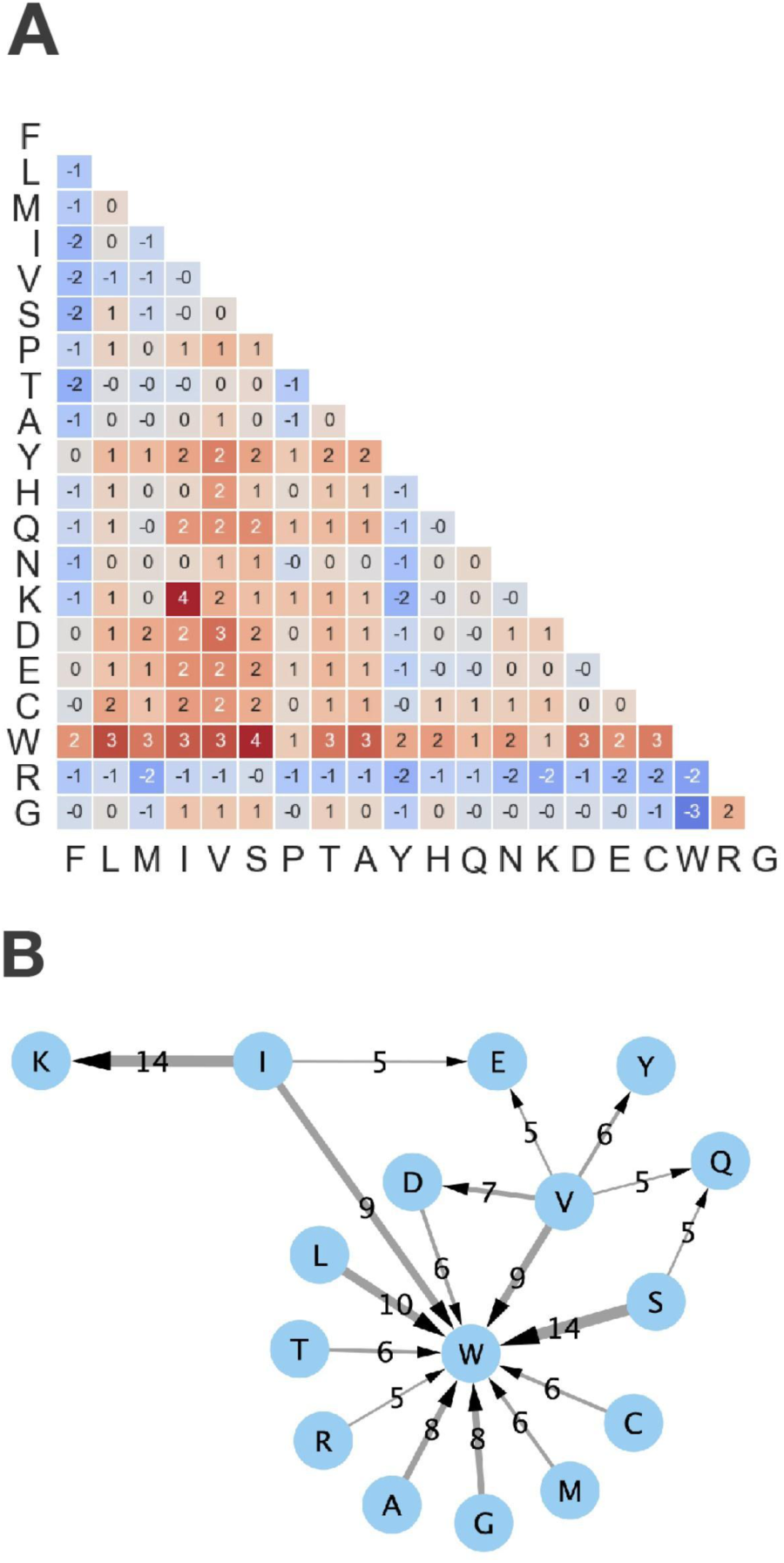
The deviation of *HA*^*3*^ from symmetry. (A) The symmetry of a substitution is measured by the log_2_ of the ratios between the probability of each substitution to the probability of its opposite substitution (resulted by swapping the source and target AAs). The probabilities are derived from a version of *HA*^*3*^ which does not include synonymous probabilities. Particularly, each row represents the conditional distribution which assigns a zero-probability to a synonymous substitution. (B) Network representation of AA substitutions that substantially deviate from symmetry (defined by ratios of at least 5, as defined in (A)). For example, serine (S) is 14 times more likely to substitute into tryptophan (W) than the other way around. Wider arrows signify stronger asymmetry (i.e., higher ratios).

### Human-specific and cross-taxa substitution matrices capture different signals

To assess the novelty of the human-specific AA substitution matrix, we compared *HA*^*3*^ with the canonical cross-taxa substitution matrices BLOSUM and PAM (Fig. 4). We observe a moderate correlation between *HA^3^* and BLOSUM_62_ (average across all 20 AAs: ρ = 0.52; Fig. 4*A*). Interestingly, BLOSUM_100_, which aims to capture much shorter evolutionary distances, differs only slightly from BLOSUM_62_ in this respect. As expected, the average correlation for BLOSUM_30_, which was designed for longer evolutionary distances, is lower (ρ=0.45). Notably, the 20 AAs exhibit high variability in this comparison. Isoleucine (I) and tryptophan (W) exhibit a higher correlation to the scoring profiles of BLOSUM_30_ than to those of BLOSUM_62_ and BLOSUM_100_.

**Figure 4.**
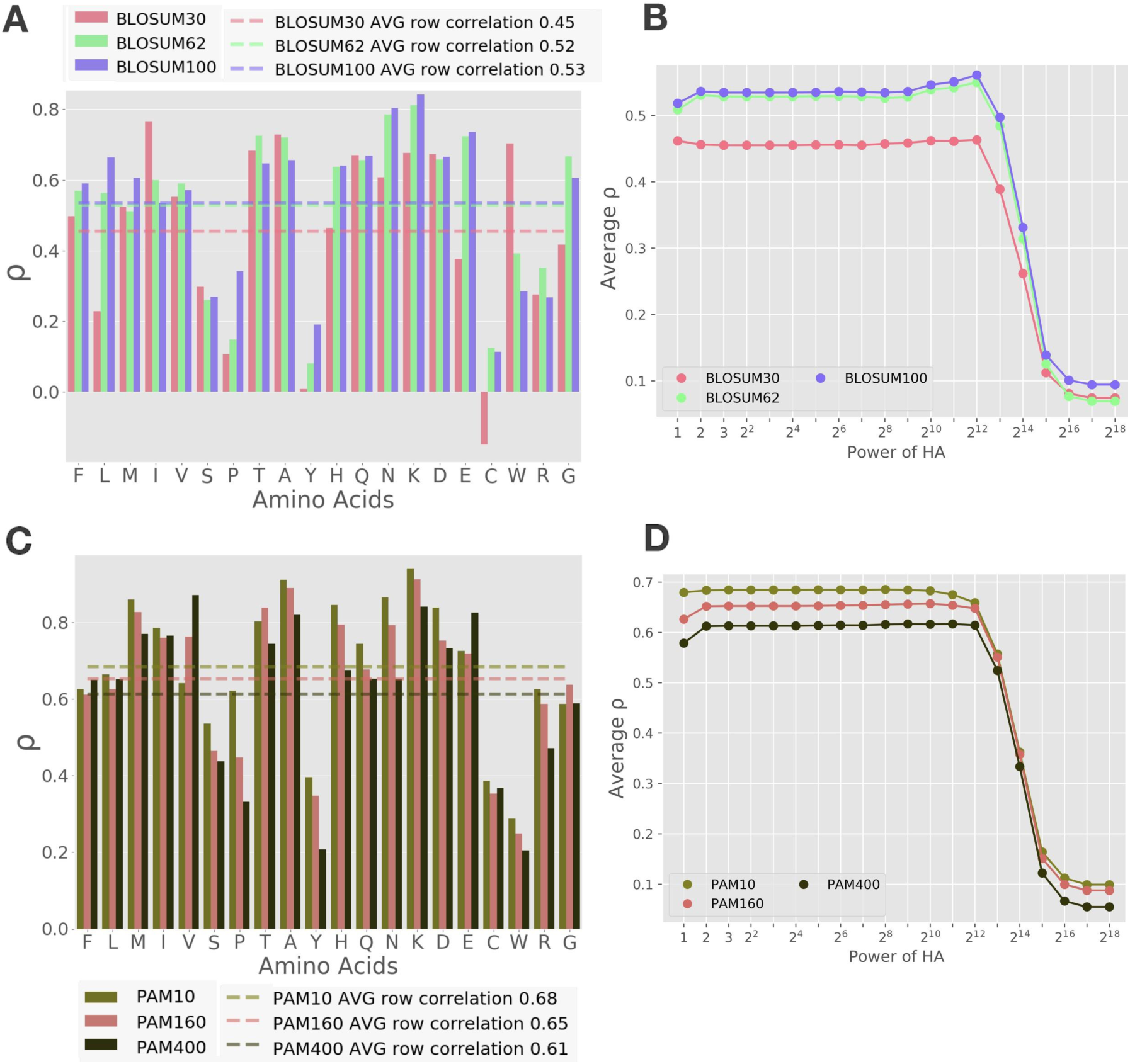
Comparison to BLOSUM and PAM substitution matrices (A) Spearman’s correlation coefficients (ρ) between the rows of *HA*^*3*^ to the corresponding rows of different versions of the BLOSUM matrices. For example, the rightmost bar measures the consistency between the substitution probabilities of Glycine (G) according to *HA*^*3*^ and the corresponding substitution scores according to BLOSUM_100_. The average correlation coefficients (across all 20 AAs) are shown as dotted lines. (B) Average Spearman’s correlation coefficients between BLOSUM matrices and different powers of *HA*^*1*^. Increasing powers of HA represent an increasing number of consecutive substitutions, thereby longer evolutionary timescales. (*C*-*D*) Same as (*A*) and (*B*), for the PAM matrices.

*HA*^*3*^ represents the expected pairwise substitution probabilities between all AAs, having gone through three consecutive single-nucleotide substitutions. It is reasonable to inquire whether the specific choice of *HA*^*3*^ substantially affects the comparison of the human-specific and cross-taxa matrices. We, therefore, consider other numbers of consecutive single-nucleotide substitutions, which can describe increasing evolutionary distances (thereby, potentially better fitting the cross-taxa matrices). We repeated the comparison between the BLOSUM matrices to *HA*^*k*^ for different values of k (Fig. 4*B*). Indeed, the observed degree of conformity is steady for a wide range of evolutionary steps (from *k*=2 to *k*=210). For higher values of k, a drop in conformity is observed, when the Markovian process reaches its stationary state. We conclude that the choice of *k*=3 is not only the minimal number of nucleotide substitutions required to represent all possible AA substitutions but is also a robust and appropriate choice.

A similar conformity analysis was performed for several PAM matrices (Fig. 4 C-D). For PAM, we observe somewhat higher correlations (average ρ of 0.61 to 0.68). The most substantial conformity of *HA^3^* is noted for PAM_10_, which represents the shortest evolutionary distance (i.e., highly similar protein sequences) that was examined. A correlation analysis was also performed for the transition matrix of PAM_1_, yielding the highest conformity (ρ=0.69, see Supplementary Figure S4). Notably, tyrosine (Y) and cysteine (C) show consistently low conformity between the human-specific matrices to all cross-taxa matrices (both BLOSUM and PAM).

### The human-specific substitution model exposes protein functional site preservation

It is natural to consider how AA substitution tendencies change in the context of protein functional sites, as these are likely to influence evolutionary pressures. To this end, we examined six prominent PTMs (acetylation, phosphorylation, N- and O-glycosylation, succinylation, and disulfide-bond) and three types of ion-binding sites (zinc, magnesium, and iron). Altogether, we examined 92,625 experimentally-validated protein site annotations (covering 12,746 unique proteins). We began by determining which AA substitutions and annotations show statistically significant changes in the context of the examined functional sites. We tested for such changes through (i) the number of variants, and (ii) their allele frequencies (AFs). All such significant substitutions are shown in Table 1 (see Supplementary Dataset S3 for non-significant results). For example, alanine (A) to threonine (T) substitutions are significantly depleted in acetylated sites with respect to both the number of observed variants (FDR *q-value* = 4.2E-10) and their allele frequencies (FDR *q-value* = 1.3E-02).

**Table 1.** Statistically significant AA substitutions in PTM sites.

| PTM annotation | From AA | To AA | # sub. w/ annotation | # sub. w/o annotation | # variants FDR q-value | AF FDR q-value | Trend* |
| --- | --- | --- | --- | --- | --- | --- | --- |
| Acetylation | A | A | 212 | 154569 | 1.94E-02 |  | E |
|  | A | T | 51 | 91640 | 4.15E-10 | 1.27E-02 | D/D |
|  | A | V | 122 | 84602 | 3.16E-02 |  | D |
|  | K | K | 398 | 48774 | 6.89E-03 |  | E |
|  | K | N | 128 | 22414 | 2.35E-02 |  | D |
|  | K | Q | 63 | 10543 |  | 4.75E-02 | D |
| Disulfide bond | C | C | 2014 | 25117 | 9.70E-32 |  | E |
|  | C | F | 311 | 5700 | 6.68E-03 |  | D |
|  | C | G | 238 | 4429 | 8.13E-03 | 2.62E-02 | D\D |
|  | C | R | 648 | 11634 | 3.93E-04 | 6.99E-05 | D\D |
|  | C | S | 412 | 8104 | 2.56E-06 | 1.01E-02 | D\D |
|  | C | W | 175 | 3205 | 4.18E-02 |  | D |
|  | C | Y | 870 | 16183 | 6.68E-08 | 5.26E-05 | D\D |
| Hydroxylation | K | K | 105 | 49091 | 7.65E-05 |  | E |
| Methylation | K | K | 54 | 49140 | 2.36E-03 |  | E |
|  | R | R | 154 | 86468 | 3.05E-02 |  | E |
| N-linked Glycosylation | N | D | 657 | 16080 | 1.79E-02 |  | D |
|  | N | H | 274 | 7345 | 6.70E-03 |  | D |
|  | N | N | 2441 | 51886 |  | 2.25E-02 | E |
| O-linked Glycosylation | S | S | 82 | 148408 |  | 1.74E-02 | E |
|  | T | T | 105 | 142197 |  | 1.44E-02 | E |
| Phosphorylation | S | A | 285 | 9066 | 4.08E-02 | 2.68E-03 | D\D |
|  | S | F | 1054 | 25354 | 4.47E-04 |  | E |
|  | S | L | 1176 | 30503 |  | 3.41E-02 | D |
|  | S | G | 706 | 21817 | 6.68E-03 |  | D |
|  | S | S | 5403 | 143087 | 1.96E-02 |  | E |
|  | S | T | 669 | 19274 |  | 3.39E-04 | D |
|  | T | T | 1471 | 140831 |  | 3.41E-02 | D |
|  | Y | H | 85 | 15654 | 1.38E-02 |  | D |
|  | Y | Y | 454 | 50960 | 3.06E-06 |  | E |
| Succinylation | K | E | 66 | 28051 |  | 2.08E-02 | D |
|  | K | K | 106 | 49066 |  | 2.08E-02 | D |
|  | K | R | 86 | 35506 |  | 2.08E-02 | E |
| Ubiquitination | K | E | 4091 | 24020 |  | 3.40E-10 | D |
|  | K | I | 324 | 2060 |  | 1.47E-03 | D |
|  | K | K | 7536 | 41630 | 1.67E-06 | 3.47E-07 | E\D |
|  | K | M | 422 | 2752 | 4.78E-02 | 5.41E-03 | D\D |
|  | K | N | 2980 | 19562 | 3.29E-10 | 7.44E-09 | D\D |
|  | K | Q | 1477 | 9129 | 4.78E-02 |  | D |
|  | K | R | 5420 | 30168 | 1.30E-03 | 2.47E-09 | E\D |
|  | K | T | 1589 | 9119 |  | 1.40E-07 | D |

\*Whether the significant differences between annotated and unannotated residues with respect to the number of variants (left) and allele frequency (AF; right) are due to enrichment (E) or depletion (D).

**Table 2.**
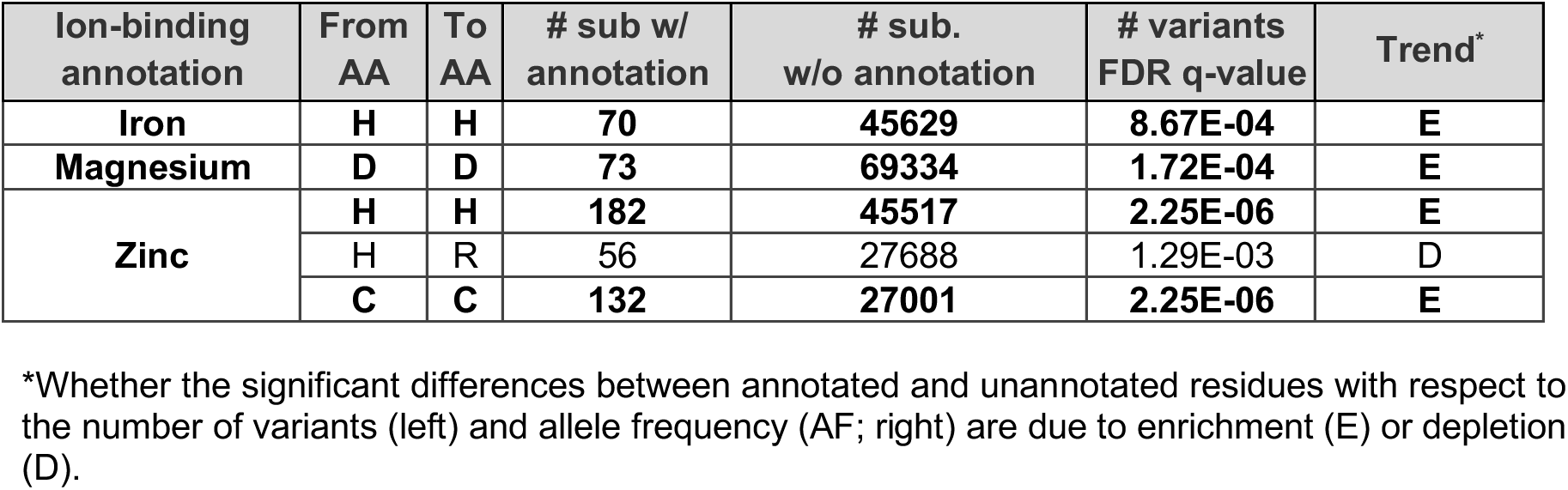
Statistically significant AA substitutions in ion-binding sites.

| Ion-binding annotation | From AA | To AA | # sub w/ annotation | # sub. w/o annotation | # variants FDR q-value | Trend* |
| --- | --- | --- | --- | --- | --- | --- |
| Iron | H | H | 70 | 45629 | 8.67E-04 | E |
| Magnesium | D | D | 73 | 69334 | 1.72E-04 | E |
| Zinc | H | H | 182 | 45517 | 2.25E-06 | E |
|  | H | R | 56 | 27688 | 1.29E-03 | D |
|  | C | C | 132 | 27001 | 2.25E-06 | E |

\*Whether the significant differences between annotated and unannotated residues with respect to the number of variants (left) and allele frequency (AF; right) are due to enrichment (E) or depletion (D).

The most significant association is the conservation of cysteine (C) residues involved in disulfide bonds (FDR *q-value* = 9.7E-32), as expected by the fundamental importance of these bonds in stabilizing the structural fold of the protein. Generally, functional residues are preserved in the majority of annotations, while nonsynonymous AA substitutions are generally depleted (Table 1). Collectively, these results expose negative selection associated with all major functional sites in the human population proteome.

Having determined which AA substitutions show significant changes in functional sites, we applied our model to examine these differences in substitution propensities (Fig. 5, Supplementary Fig. S4). Specifically, we considered the entry-wise ratios of *HA*^*3*^ within annotated and unannotated sites by reconstructing the matrix for each of the two conditions. AA substitutions that appear particularly unfavored in specific functional contexts are acetylated lysine (K) to glutamine (Q), zinc-binding histidine (H) to arginine (R), and hydroxylated alanine (A) to valine (V) or threonine (T). Lysine (K) plays an essential role under many PTMs, several of which exhibit strong negative selection under our model. The majority of succinyl-modified lysine residues overlap with acetylation (22). lysine modified by ubiquitin, acetyl, and succinyl presents a lower tendency to change to other AAs, compared to an unmodified lysine. We did not analyze additional types of lysine PTMs (sumoylation, methylation) due to a lack of experimental evidence. We conclude that a human-centric model of AA substitution propensities is a powerful and intuitive tool to study a variety of short-term evolutionary contexts, including PTMs and the amino acids that support and coordinate ion-binding sites.

**Figure 5.**
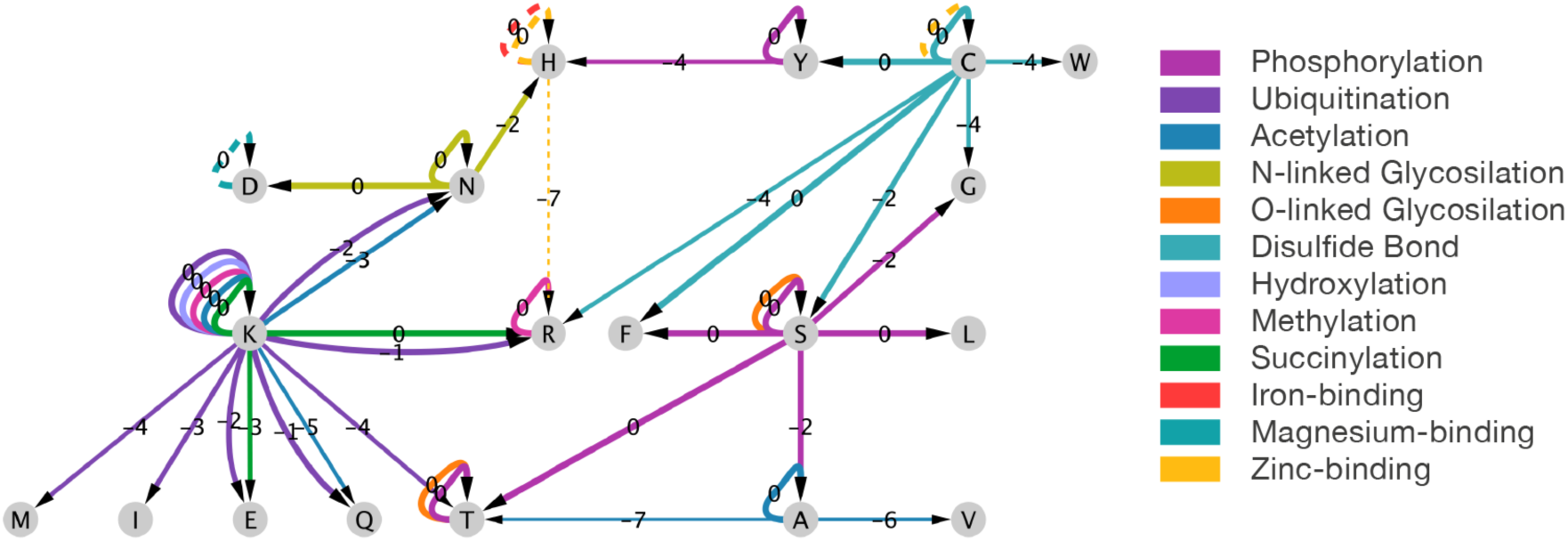
Network representation of AA substitutions exhibiting significantly different tendencies in PTM and ion-binding sites. Each edge represents a substitution between a source to a target AA in the context of a specific annotation (indicated by edge color). The edges are annotated with the log_2_ ratio of the substitution probabilities between sites with and without the annotation. Arrow widths signify the magnitude of these ratios. AAs that allow annotations, and are therefore potential substitution sources, are marked as orange squares.

## Discussion

In this study, we have presented a set of novel, data-driven codon and AA substitution matrices, which is based on the natural occurrence of genetic variations in the healthy human population. This modern-human centric approach implies an evolutionary timescale of up to tens of thousands of years, as opposed to the classic BLOSUM and PAM matrices, which are based on proteomic sequences from organisms whose common ancestor goes back hundreds of millions of years (23). Despite the different order of magnitude of evolutionary timescales, *HA*^*3*^ still shows a moderate-to-high correlation with the BLOSUM and PAM matrices (Fig. 4). Moreover, the matrices that reflect shorter evolutionary distances (e.g., BLOSUM_100_ and PAM_100_) exhibit higher similarity to the human-centric matrix. On the other hand, we do observe that some amino-acids, most notably cysteine (C), tyrosine (Y) and tryptophan (W), seem to behave very differently in the human context. These amino acids specify unique biochemical features which are influential on protein structure properties (e.g., tyrosine has the lowest frequency for alpha-helix), function (24), and explicitly on protein interactions. Specifically, C, Y, W have strong interface propensity (25) and negative solvation potential (26). Interestingly, these amino-acids are implicated with the highest probability for causing disease in human de-novo rare mutations (27).

We argue that a human-centric model that reflects short-term evolution is more appropriate for accurate assessment of the impact of human mutations in coding genes, which is essential for human genetic consulting and identifying causal mutations in rare diseases. For this task, many prediction tools and algorithms have been developed (e.g. SIFT, PolyPhen2, CADD, SNAP2, FIRM (28–32)). Many of these tools incorporate long-range evolutionary information from cross-species conservation into the underlying model (33, 34). We anticipate that the incorporation of short-term human baseline models such as *HA*^*3*^ could be beneficial to these efforts.

Interestingly, even the very short-term view, explored in this study, exposes a clear and robust signal of negative natural selection at codon and amino-acid resolution (Fig. 3*A*). Unsurprisingly, synonymous substitutions (the diagonal values of the matrix) exhibit higher propensities than would be expected by the nucleotide compositions of the codons, while missense substitutions are suppressed. Our model further exposes notable differences between substitutions, with some showing an order-of-magnitude stronger selection than others. While most AAs tend to substitute into tryptophan (W) more than the other way around, the AA with the most extreme tendency is Serine (S), where the ratio of propensities is 14. Another extreme case is the substitution of isoleucine (I) to lysine (K), which is ~14 times less frequent than would be expected by the codons of these amino-acids. Examining the same substitution in codon resolution, we reveal that a single codon substitution primarily drives this signal from ATA (I) to AAA (K) (Supplementary Fig. S1, Supplementary Dataset S2).

Having observed a strong evolutionary signal in our model of amino-acid substitutions, we sought to consider its implication on protein function, specifically in the case of PTMs and ion-binding residues (Table 1 and Fig. 5). At a cellular level, key processes such as apoptosis, cell division, signal transduction, and cellular communication are governed by a network of PTMs (35). In humans, most proteins are subject to PTMs, and the combinatorics of PTMs greatly increase the proteome’s functional repertoire (36, 37). While we discuss each of the PTMs separately (Fig. 5), it is known that different types of PTMs may occur on the same residue (38). For example, lysine (K) participates in many of the major PTM types, with over half of the acetylation and a third of ubiquitination co-occur at overlapping lysine (K) (39).

Although ~200 types of PTMs have been detected by mass spectrometry (MS) (40–42), only a handful were broadly used and systematically studied (43, 44). Whether PTM sites are under neutral, negative, or positive selection in humans remained an unsolved question (3, 45, 46). In this study, we report a strong negative selection signal that is evident for many of the PTM types and ion-binding sites. Recall that PTMs are highly volatile and condition-specific, and their annotations are sensitive to experimental protocols (47, 48). Despite these notable sources of noise (40, 49), we detected several statistically significant substitutions that are enriched or depleted within specific PTMs. Particularly salient are functional contexts with clinical implications such as cysteine disulfide bonds (50), phosphotyrosine (relative to phosphoserine and phosphothreonine) (51) and ion-binding sites (52). Many rare monogenic diseases result from point mutations in codons of cysteine, which destroy essential disulfide bridges bonds (50). As many of the discussed PTMs are often positioned at protein tails (e.g., N’-acetylation), loops and disordered regions (e.g., phosphorylation, glycosylation, and ubiquitination), the negative selection of PTMs exposed in this work provides evidence for the functional importance of these protein regions. Regions of low sequence similarity across protein homologs are restricted to protein tails, loops, and interdomain linkers. Based on the low cross-species conservation of such regions, we anticipate that the functional PTM conservation signal that is exposed in this study has already faded in the context of a long-range cross-taxa evolution.

A model with simple probabilistic interpretability was a key consideration in the design of the substitution matrices presented in this work. Each row in our matrices represents the estimated distribution of conditional substitution probabilities for a given amino-acid, and can, therefore, be observed and interpreted independently. In other words, each of our matrices represents a Markov chain, which is a natural way to model substitutions (e.g., the PAM matrices), since it holds a convenient property by which more distant substitutions involving multiple steps of the Markov chain can be easily derived from the baseline single-step matrix. In this manner, we extrapolate *HC*^*3*^ and *HA*^*3*^ from the data-driven matrices *HC*^*1*^ and *HA*^*1*^, respectively. We also considered higher numbers of consecutive substitutions (Fig. 4*B*, 4*D*) and observed that the described Markov chain is robust for a broad range of timescales. On the other hand, it would be difficult to make predictions about long-term evolutionary distances that are based on short-term dynamics alone (53). For tasks that involve long evolutionary distances across species, the PAM and BLOSUM matrices are likely to be more suitable.

Another essential property of the constructed matrices we present is their asymmetry (Fig. 4). Conveying directionality in our model was made possible thanks to our reliance on genetic variants that are associated with allele frequencies (for further details, see Methods). In contrast, the BLOSUM matrices assume symmetry of substitutions by design, since they were constructed from cross-species multiple sequence alignment data and are transformed into scores using a log-odds ratio. The symmetric PAM scoring matrices are derived from asymmetric transition matrices and are symmetric as a result of the transformation into a scoring model. Having considered the directionality of amino-acid and codon substitution in the human population, we have found a substantial observed asymmetry for some amino acids (Fig. 3*B*). Specifically, tryptophan (W) and valine (V) appear to be central hubs of such directional tendencies. Notably, specific AAs can be signified by being targets (tend to be substituted to more than by) or sources (vice versa). We found that tryptophan (W), and to a lesser extent, glutamic acid (E) and glutamine (Q), are attractors. Valine (V), and to a lesser extent, serine (S) and isoleucine (I), act as sources. Repeating the analysis (as in Fig. 3*B*) using a relaxed threshold (Supplementary Fig. S3) substantiates the target/source AA partition. Interestingly, the AAs marked as source include all AAs with 6 codons (R, S, L) and most AAs with 4 codons. Our methodology allows us to investigate these insights also at the resolution of codons, exposing that patterns of asymmetry vary between codons of the same AAs (Supplementary Fig. S2). For example, the asymmetric substitution of isoleucine (I) to lysine (K) is dominated by a specific codon (ATA substitution to AAA). In this view, a particular case is of serine (S) codons. The 6 codons are may be thought of as composed of two sets of codons, which exhibit distinct substitution patterns, with broad functional implications (17). Specifically, regions that are subjected to an accelerated evolution tend to substitute within one serine codon set, while the conserved region in the same proteins tend to substitute within the other serine codon set. Our findings support the notion that in proteins, different substitution paths are likely to have functional implications (17).

In summary, we have constructed human-specific substitution matrices and characterized their unique properties. Given the robustness and interpretability of these matrices, we encourage their use as a baseline model for codon and amino-acid substitutions in the healthy human population. The demonstrated PTM and ion-binding analyses illustrated in this work is an example of the type of studies that could be performed given a reliable background model. We anticipate that more genetic and proteomic functional elements can be exposed through these matrices. To allow the extension of this methodology to other datasets, including specific human subpopulations, we provide the source code of our methods (see Methods). We anticipate that such a methodology could be easily applied to other organisms with multiple genetic samples (e.g., mouse (54)). Viral isolates, which often contain thousands of samples per strain, could be a particularly relevant subject for such methods.

## Materials and Methods

### Data

To construct human-specific codon and AA substitution matrices, we combined genetic variation data with functional annotations of coding genes (Fig. 1*A*). Human genetic variations were extracted from ExAC (18), which provides a good tradeoff of quantity and quality of genetic data in the human population (European origin biased). Importantly, the cohort of ExAC was chosen to minimize bias of pathogenic variants, and exclude individuals with rare genetic diseases. We treat each variant as a substitution from the major allele, which we defined as the most frequent allele in the population, to the minor allele, which is any other allele observed in the specific exomic location.

Functional gene annotations were taken from the UniProt database and the GENCODE project. The exact procedure of combining genetic and proteomic annotations is described in (28). Briefly, we used version 19 of GENCODE (compatible with version GRCh37 of the human reference genome, which was used by ExAC). We recovered the DNA sequences of the genes annotated in GENCODE using UCSC’s reference genome. We considered only the protein-coding regions of genes (annotated as “CDS” in GENCODE). Protein sequences and protein annotations (e.g., PTMs; see next section) were taken from UniProt for all 20,168 reviewed human proteins (from the SwissProt section). We only considered the GENCODE gene isoforms identical to the primary UniProt protein sequences. We discarded genes that failed this exact one-to-one mapping, ending up with 18,115 successfully mapped genes. These combined genetic-proteomic gene entities allowed us to determine the protein-level consequences of genetic variants (e.g., synonymous, missense, or nonsense). This pipeline is available as an independent open-source Python library (https://github.com/nadavbra/geneffect).

ExAC identified 7,404,909 high-quality human genomic variants. From this dataset, indels were removed, and 7,087,528 SNVs remained. 326,369 genomic positions contained multiallelic variants, which were counted as 631,985 nucleotide substitutions, contributing 305,616 additional variants. 83,045 nonsense mutation variants were discarded. Of the 7,627,480 nucleotide substitutions, 4,786,729 were within the coding regions of the 18,115 mapped protein-coding genes. These synonymous and missense SNVs comprised the final dataset used in this work.

### PTMs and ion-binding site annotations

PTMs and ion-binding site annotations were extracted from UniProt, except for ubiquitination annotations, which were obtained from PhosphoSitePlus (55). For a small number of cases, PTM proteomic locations of variants mapped to locations on the reference genome that code for inappropriate AAs. Importantly, both UniProt and Phosphosite annotations are supported by strong experimental evidence and literature.

### Constructing probabilistic human-specific substitution matrices

*HC*^*1*^ represents the estimated substitution probability of each pair of the 61 coding codons (i.e., excluding the three stop codons). Each non-diagonal entry of *HC*^*1*^ was calculated by:

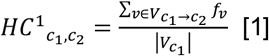

Where 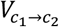 denotes the set of all variants substituting codon *c*_1_ to *c*_2_ (*c*_1_ ≠ *c*_2_), 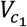 the set of all variants substituting *c*_1_ to any codon (including a self-substitution), and *f*_*ν*_ the frequency of the *c*_2_ allele in a variant *ν* substituting *c*_1_ to *c*_2_ (as derived from ExAC).

The diagonal values were calculated by:

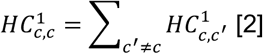

We assumed the directionality of each variant to be from the major to the minor allele. In particular, we always have that *f*_*ν*_ ≤ 0.5. Note that the major allele is not always the reference allele (the allele matching the human reference genome). This is not the case in ~8% of the ~5M processed variants. Since the aggregated data offers information about each exomic position independently, it essentially ignores the effect of dependence between variants.

The non-substitution (diagonal) values of the matrix complement the sum of each row to 1. This, along with the normalization in Eq.1, yields a row-stochastic matrix, meaning that each row *i* of the matrix can be interpreted as the distribution of conditional substitution probabilities (from the *c*_*i*_ codon to every other codon). Each entry, in turn, may be interpreted as the conditional probability of observing *c*_2_ in a genomic position where *c*_1_ is the major allele.

The amino-acid substitution matrix *HA*^*1*^ is derived from *HC*^*1*^ by considering codon frequencies:

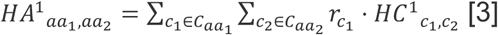

Where 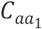 and 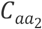 denote the sets of codons that code for the amino-acids *aa*_1_ and *aa*_2_, respectively, and 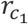 is the frequency in which *aa*_1_ is coded by the *c*_1_ codon, relative to all the codons of *aa*_1_ (i.e.,., 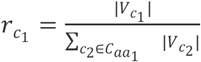).

As with *HC^1^*, an entry of *HA^1^* may be conceived as the conditional probability of observing amino-acid *aa*_2_ in a proteomic position in which *aa*_1_ is the major allele.

As *HC*^*1*^ and *HA*^*1*^ were generated through statistical analysis of single-nucleotide variations, substitutions between codons that differ in more than one nucleotide cannot be directly inferred. Since most codon pairs differ by more than one nucleotide, both *HC*^*1*^ and *HA*^*1*^ are sparse matrices. To estimate substitution frequencies between pairs of codons requiring multiple consecutive substitution events to transition between them, we treat *HC*^*1*^ and *HA*^*1*^ as transition matrices of Markov chains. For every number of consecutive substitutions *k*, the *k*-th power matrices *HC*^*k*^ and *HA*^*k*^ represent the Markov chains that are a result of repeating the original Markov chains *k* times. To obtain a complete substitution matrix (i.e., with non-zero substitution probabilities for all possible substitutions), we chose to take *HA*^*1*^ to the power of 3. This is *HA*^*3*^, since 3 is the lowest number of nucleotide substitutions required between each pair of coding-codons. This is a desired quality, since we aim to model substitution propensities between closely related sequences which require a low number of evolutionary steps. As demonstrated (Fig. 4), this choice has little impact on our analysis.

### Comparing the amino-acid substitution model to a nucleotide substitution model

To examine to what extent our substitution matrix reflects evolutionary signals at the amino-acid level, we compared it to a matrix derived from nucleotide substitution frequencies, which we used as a simple background model (Fig. 2*B*). To this end, we first constructed a 4×4 nucleotide substitution matrix (Fig. 2*A*), derived from the same set of variants used to construct *HC*^*1*^. We then used this nucleotide-level matrix to derive an expected 61×61 codon substitution matrix, by considering the probability of a codon substitution to be the product of the probabilities of the single-nucleotide substitutions involved in that codon (e.g., the substitution of CTG to TTA was assigned the probability of C to T multiplied by the probability of T to T and the probability of G to A). This codon-level substitution matrix was then projected into a 20×20 AA substitution matrix, through the same process used to convert *HC*^*1*^ to *HA*^*1*^ (see previous section). The resulting AA-level matrix reflects the expected probabilities of AA substitutions given only the substitution preferences of single nucleotides while assuming a lack of evolutionary pressure at the codon, or amino-acid, level. By dividing the empirical AA substitution matrix, *HA*^*1*^, with that background model (entry-wise), we obtained the observed-to-expected probability ratios of all the AA substitutions and could determine which AA substitutions show a significant deviation from that background model.

### Deviation from symmetry

To quantify the strength of directionality observed in the substitution matrix, we measured the deviation of amino-acid substitutions from symmetry by calculating the ratios between the conditional probabilities of substitutions to the conditional probabilities of the opposite substitutions. For example, the directionality of the substitution of lysine (K) to arginine (R) under *HA*^*3*^ is measured by 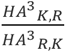. In other words, we divided *HA*^*3*^, entry-wise, by (*HA*^*3*^)^T^. To enhance readability, the matrix shown in Fig. 4*A* is lower triangular, and its entries are transformed by log_2_. This means that positive entries signify substitutions that are preferred over their opposite substitution, and negative entries signify the opposite.

### Comparison to BLOSUM and PAM

To compare our substitution models to the score matrices of BLOSUM, Spearman’s correlation coefficient (ρ) was measured for each row of *HA*^*3*^ with each corresponding row of the examined BLOSUM matrix. For instance, ρ was calculated for the arginine (R) row of *HA*^*3*^ and the arginine (R) row of BLOSUM_100_. This process was done for all rows of *HA^3^* against their corresponding rows in each BLOSUM version (Fig 4*A*). For each BLOSUM version, its average correlation with *HA*^*3*^ was used to summarize these measurements. Furthermore, to examine the effect of the power of the *HA*^*1*^ matrix to the correlation to BLOSUM, the average correlation coefficient was calculated for each BLOSUM version, and different powers of *HA*^*1*^ (Fig 4*B*). These analyses were repeated for the PAM matrices (Fig 4*C*-*D*).

### Functional annotation analysis

To demonstrate the capacity of our human-specific substitution model to reflect protein functional annotations (Table 1 and Fig. 5), we examined nine major PTMs (acetylation, hydroxylation, disulfide bond formation, methylation, N-linked glycosylation, O-linked glycosylation, phosphorylation, succinylation and ubiquitination) and three types of ion-binding sites (zinc, magnesium and iron). For each PTM or ion-binding site, we considered the set of variants at the annotated proteomic locations, and generated codon and amino-acid substitution matrices corresponding to that subset of variants (e.g. *HA*^*1*^_iron-binding_, *HC*^*3*^_ubiquitination_, etc.). These matrices were generated through the same method by which the global matrices (e.g. *HA*^*1*^, *HC*^*3*^) were constructed, and they differ only by the subset of used variants (within ExAC dataset). To highlight the unique aspects of the substitution profiles for these functionally annotated sites, compared to unannotated sites (Fig. 5), each annotation-specific matrix was divided by its corresponding non-annotation matrix, element-wise (i.e. the matrix derived from all the other variants).

To test the significance of each annotation-specific substitution (e.g., lysine (K) to proline (P) in ubiquitination sites), we examined two complementary aspects of significance, based on either i) the number of annotated variants or ii) their allele frequencies (AF). In terms of the number of variants, a significant substitution may exhibit a significantly higher or lower number of variants in sites annotated by that PTM\ion-binding. To test whether a substitution *aa*_1_ → *aa*_2_ is significantly associated with an annotation in terms of the number of variants, we considered the set of all variants whose major allele is *aa*_1_ and used Fisher’s exact test to determine if the subset of these variants that substitute into *aa*_2_ is enriched with the subset of variants with that annotation. Likewise, to test differences in AF, we used Mann-Whitney U test (two-sided) to compare the AF of the variants of the tested substitution which are in annotated vs. unannotated sites (e.g. lysine (K) to proline (P) variants in ubiquitinated vs. non-ubiquitinated sites). In both tests, we required a sample size of at least 50 annotated variants. To control the false discovery rate, Benjamini– Hochberg FDR was applied for each of the two types of tests, across all annotation-specific substitutions, with a significance threshold of 0.05. In this work we show the annotation-specific substitutions that are FDR-significant according to at least one of the two tests (Table 1 and Fig. 5). Significant annotation-specific substitutions are labeled as either enriched (E) or depleted (D) for each of the two tests. With respect to the number of variants, we consider it to be enriched if the odds-ratio is greater than 1. With respect to the AF test, we consider it to be enriched when the average AF of annotated variants is greater than that of unannotated variants.

## Data and code availability

The data that support the findings of this study can be downloaded from ftp.broadinstitute.org/pub/ExAC_release/release0.3/ExAC.r0.3.sites.vep.vcf.gz. Matrices, codon counts, and further results are available in the supplementary data. Relevant code generated during this study is available at github.com/tairsha/taxa-specific-substitution-matrix.

## Acknowledgments

We thank Nathan Linial (The Hebrew University of Jerusalem) for his support and suggestions throughout this research. We thank Liran Carmel (The Hebrew University of Jerusalem) and Uri Hirshberg (Haifa University) for their advice and useful comments. T.S. fellowship was partially funded by Yad Hanadiv grant (#9660/ 2019).

