## Supplemental Materials for "Evolutionary and Functional Lessons from Human-Specific Amino-Acid Substitution Matrices"

Supplementary Information for

### Evolutionary and Functional Lessons from Human-Specific Amino-Acid Substitution Matrices Paste the full author list here

Tair Shauli1, Nadav Brandes1, Michal Linial 2

1The Rachel and Selim Benin School of Computer Science and Engineering, 2Department of Biological Chemistry, Institute of Life Sciences, The Hebrew University of Jerusalem, Jerusalem, Israel

Corresponding Author: Michal Linial

Figures S1 to S4
Legends for Datasets S1 to S4

**This PDF file includes:**

**Other supplementary materials for this manuscript include the following:**

Datasets S1 to S4

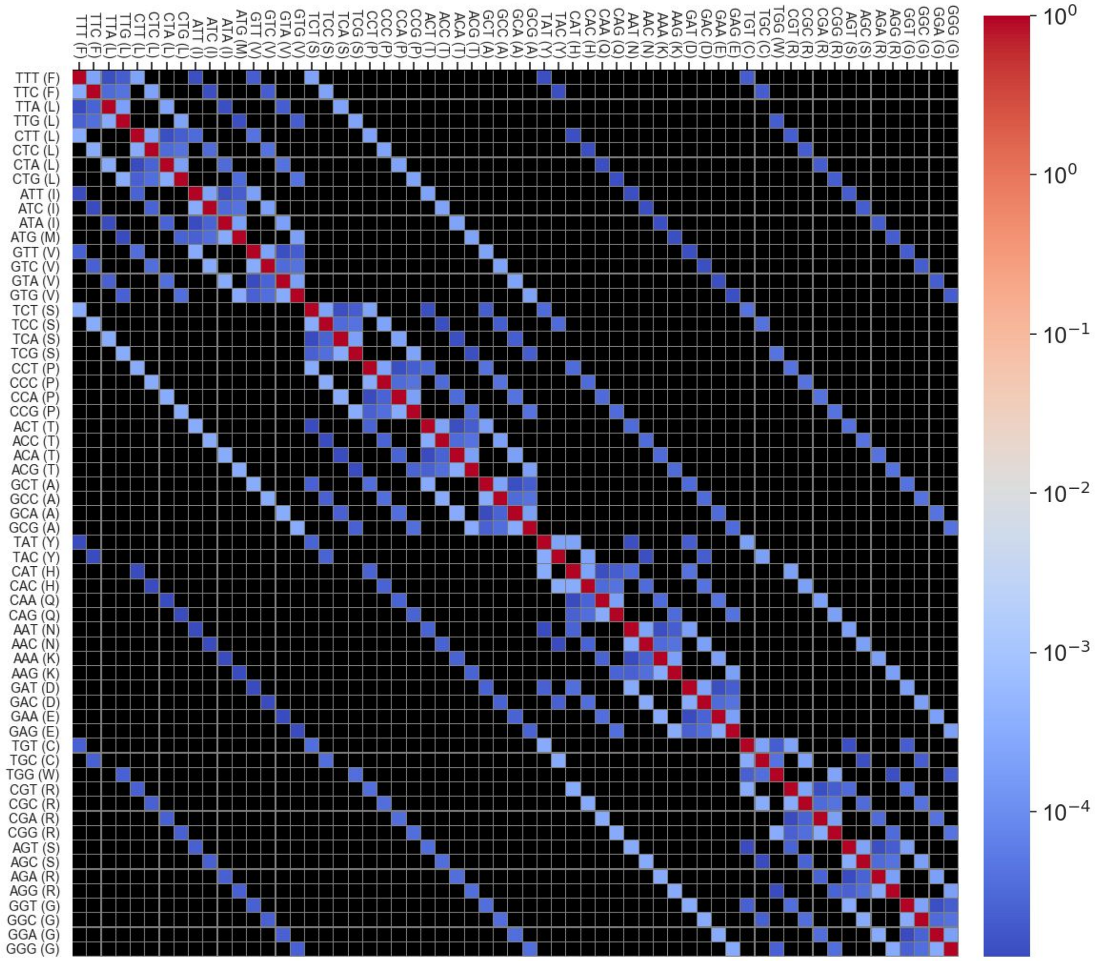


**Fig. S1.** The values of $HC^1$.

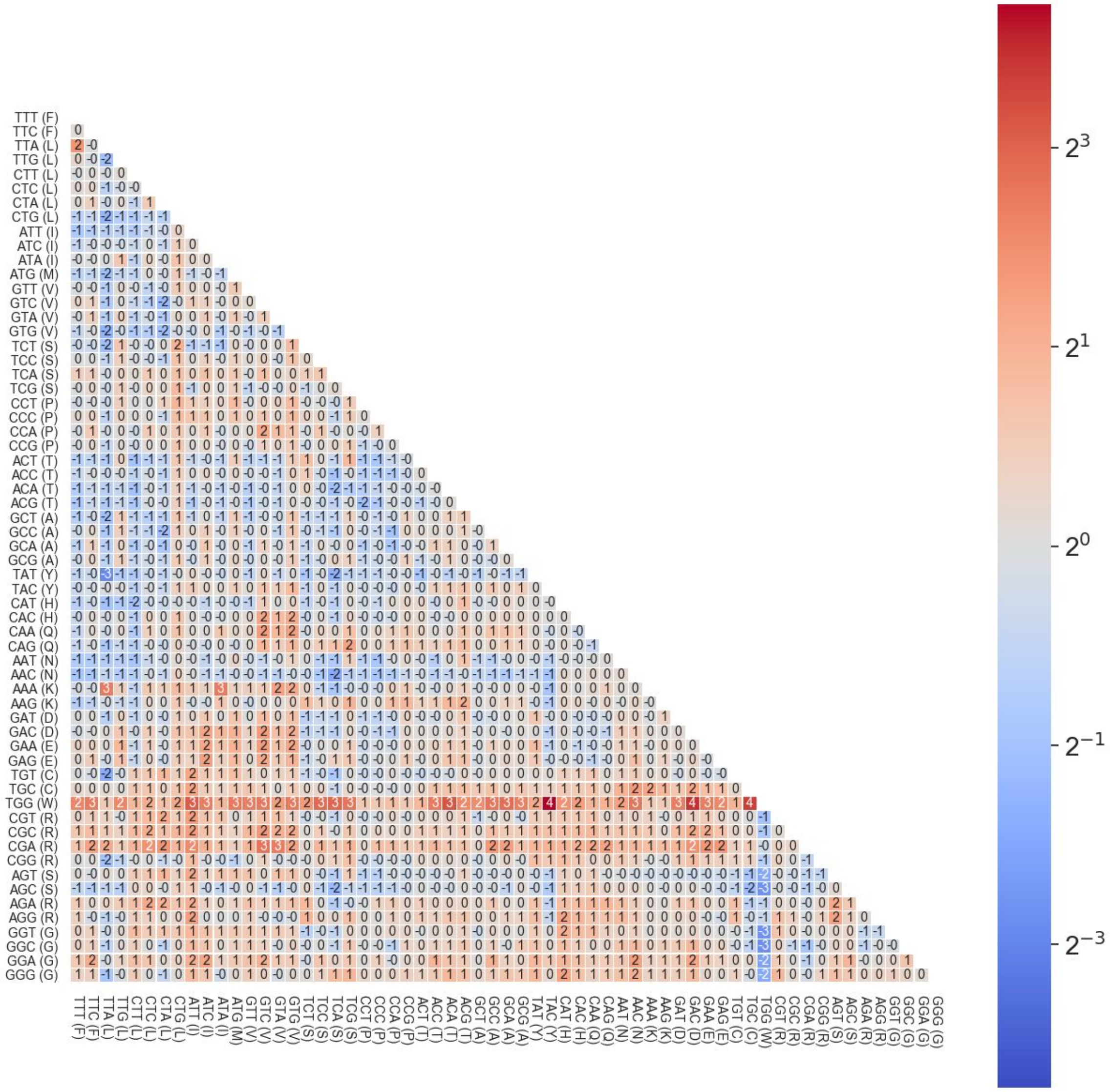


**Fig. S2.** The deviation of $HC^3$ from symmetry. The symmetry of a substitution is measured by the $\log_2$ of the ratios between the probability of each substitution to the probability of its opposite substitution (resulted by swapping the source and target codons). The probabilities are derived from a version of $HC^3$ which does not include synonymous probabilities. Particularly, each row represents the conditional distribution which assigns a zero-probability to a synonymous substitution.

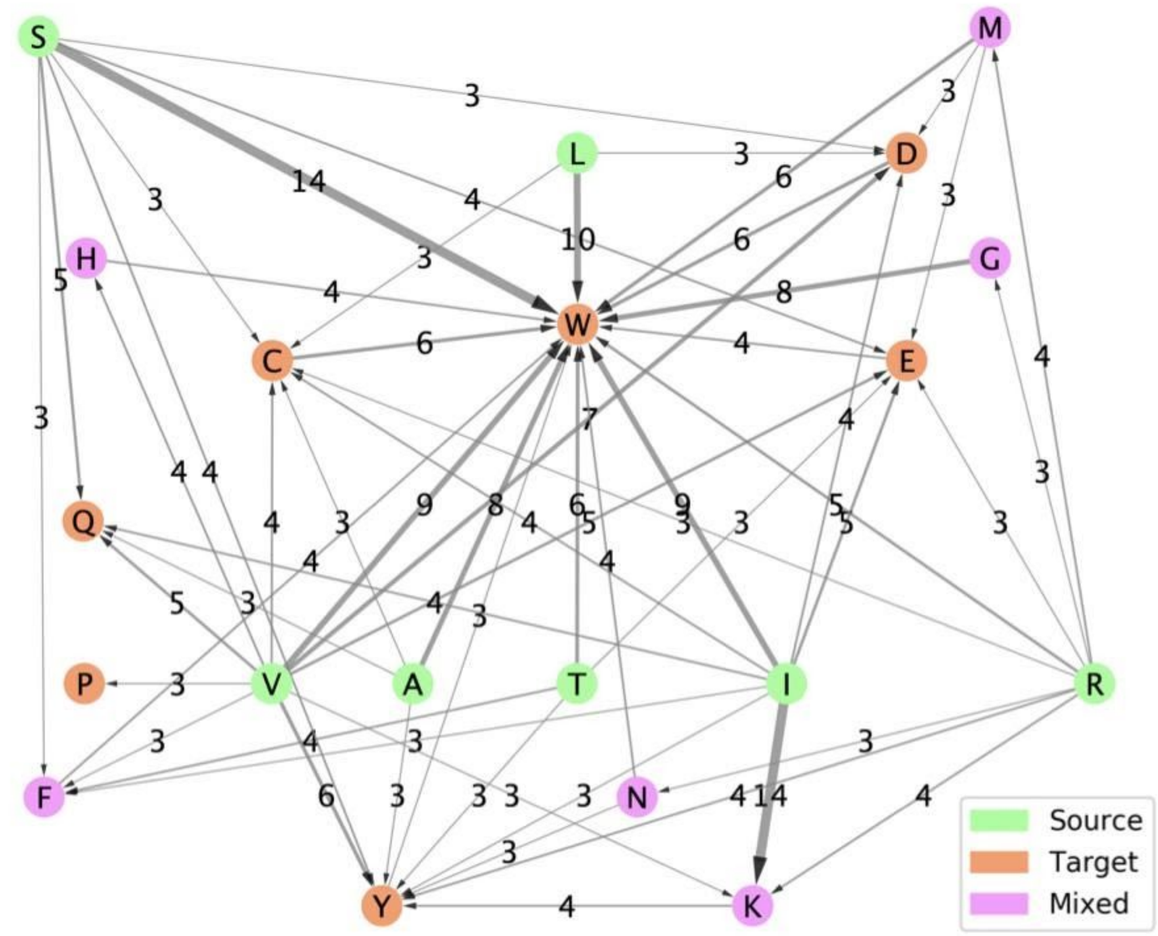


**Fig. S3.** Network representation of amino-acid (AA) substitutions that substantially deviate from symmetry (defined by ratios of at least 3, as defined in Fig.3A). For example, serine (S) is 14 times more likely to substitute into tryptophan (W) than the other way around. Wider arrows signify stronger asymmetry (i.e., higher ratios). Ratios are rounded. Colors specify whether an AA is a source in ≥ 80% of the substitutions, of a target in ≥ 80% of the substitutions, or nether.

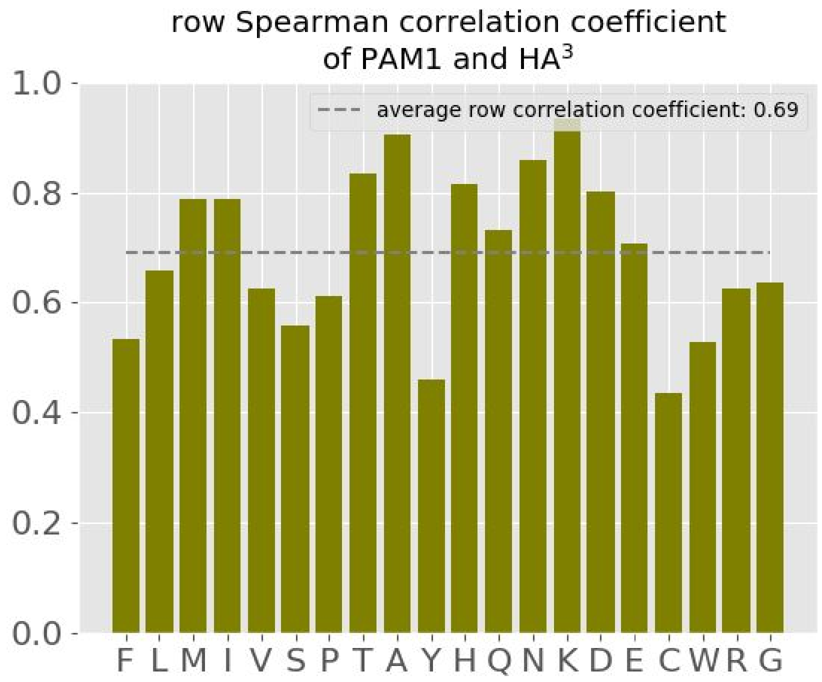


**Fig. S4.** Comparison of HA3 to $PAM_1$. Spearman's correlation coefficient (ρ) between the rows of HA3 to the corresponding rows of $PAM_1$. The average correlation coefficient (across all 20 AAs) is shown as a dotted line.

**Dataset S1 (separate file).** Full numeric values of $HC^1$, $HC^3$, $HA^1$, $HA^3$.

**Dataset S2 (separate file).** Full numeric values of $HN^1$, $HCN^1$, $HAN^1$, $HAN^3$.

**Dataset S3 (separate file).** Complete results for the statistical tests performed for each type of post-translational modification and ion-binding site annotation.

**Dataset S4 (separate file).** Full numeric values of $HA^1$ as calculated for each post-translational modification and ion-binding site annotation.
